# Visceral Fat Metabolic Indices and Thyroid Function: A Stratified Analysis of Non-Linear Associations and Population Modifiers Using NHANES Data

**DOI:** 10.64898/2026.03.19.713076

**Authors:** Kejun Chen, Wu Pan, Zhujun Yang

## Abstract

**Background:** This study was designed to investigate the relationship between visceral fat metabolic score (METS-VF), lipid accumulation product (LAP), visceral adiposity index (VAI) and thyroid function.

**Methods:** Utilizing data from the National Health and Nutrition Examination Survey (NHANES) 2007-2012, participants were excluded if they lacked data on METS-VF, LAP, VAI or thyroid function, or were under 18 years of age. Multiple linear regression, smooth curve fitting, and subgroup analyses were performed to determine the independent relationship between lipid accumulation and thyroid function.

**Results:** After full covariate adjustment, all three visceral adiposity indices showed significant positive associations with FT3 (LAP: β=0.028, VAI: β=0.024, METS-VF: β=0.026; all P<0.001), FT3/FT4 ratio, TT3, TT4, and TgAb. LAP and VAI demonstrated inverse associations with FT4 (β=-0.218 and −0.183, respectively; both P<0.001), while VAI and METS-VF were positively associated with TSH (β=0.149, P=0.041; β=0.167, P=0.025). Quartile analyses confirmed dose-dependent relationships, with Q4 participants showing elevated FT3, FT3/FT4, TT3, TT4, and reduced FT4 compared to Q1. RCS analyses revealed distinct non-linear patterns: LAP exhibited non-linearity with FT3, TSH, TT3, and TT4 (all P-nonlinear<0.05) but linear inverse associations with FT4. VAI displayed reverse L-shaped curves for FT3, TSH, and TT3 with plateaus at higher levels, while TT4 showed an inverted U-shape. METS-VF demonstrated non-linear increases for FT3 and TT3, linear associations with TSH and TT4, and an inverted U-curve for FT4. Stratified analyses identified age, race, and smoking as consistent modifiers of FT3/FT4 associations across all indices (interaction P<0.05), with stronger effects in younger/older adults, males, White participants, and high-income groups. TT3 and TT4 modification patterns varied by index. Thyroid autoantibodies showed minimal associations across all indices.

**Conclusion:** Visceral lipid accumulation is closely associated with thyroid dysfunction, and this association exhibits significant non-linear characteristics, which are modulated by factors such as age, race, and lifestyle habits. These findings provide new perspectives for the early identification and intervention of obesity-related thyroid dysfunction.

## 1 Introduction

The thyroid gland, recognized as the largest endocrine organ in the human body, is critically involved in regulating metabolism, maintaining growth and development, and supporting cardiovascular function^1^. Two primary hormones are secreted by the thyroid: thyroxine (T4) and triiodothyronine (T3), with T4 serving as a prohormone that is converted to the more biologically active T3 through deiodinase activity in peripheral tissues^2^. Thyroid hormone levels are maintained through precise regulation by the hypothalamic-pituitary-thyroid (HPT) axis, which forms a complete negative feedback loop whereby elevated serum free thyroid hormone concentrations inhibit the secretion of thyrotropin-releasing hormone (TRH) from the hypothalamus and thyroid-stimulating hormone (TSH) from the pituitary gland, thereby modulating thyroid hormone synthesis and release^3^. Thyroid dysfunction represents a common endocrine disorder, encompassing hyperthyroidism (prevalence approximately 0.5%-2%) and hypothyroidism (prevalence approximately 1%-5%), with global incidence rates showing an upward trend^4^. Even subclinical thyroid dysfunction has been associated with increased risks of various metabolic disorders and cardiovascular diseases^5^.

Obesity has emerged as a major public health concern in the 21st century, significantly increasing the risk of diabetes, cardiovascular disease, and certain cancers^6^. It is projected that by 2030, the total number of adults affected by high BMI in China will reach 515 million, ranking first globally^7^. However, traditional obesity assessment indicators such as BMI have fundamental limitations: they cannot distinguish between muscle and adipose tissue, and critically, they fail to account for visceral fat distribution^8^. Compared to subcutaneous fat, visceral adipose tissue is more metabolically active, secreting numerous pro-inflammatory factors that are closely associated with insulin resistance, vascular endothelial dysfunction, and atherosclerosis^9^.

To overcome the limitations of traditional indicators, multiple visceral fat accumulation assessment indices have been developed in recent years. The Metabolic Score for Visceral Fat (METS-VF) integrates parameters including the metabolic score for insulin resistance, waist-to-height ratio, age, and sex, demonstrating good correlation with visceral fat area measured by dual-energy X-ray absorptiometry (r=0.77) and achieving area under the curve values of 0.92-0.94 for predicting visceral obesity^10^. The Lipid Accumulation Product (LAP), calculated based on waist circumference and triglycerides, reflects visceral fat functional status and shows superior correlation with insulin sensitivity and metabolic syndrome compared to BMI^11^. The Visceral Adiposity Index (VAI) is a sex-specific comprehensive indicator that combines BMI, waist circumference, triglycerides, and high-density lipoprotein cholesterol parameters, enabling indirect assessment of visceral fat distribution and function^12^. These indices offer practical advantages as they are non-invasive, cost-effective, and readily calculable from routine clinical measurements.

Thyroid hormones play a central role in energy metabolism through multiple mechanisms that regulate lipid synthesis, breakdown, and transport^13^. T3 activates thermogenic functions in brown adipose tissue, promotes fatty acid oxidation, and increases energy expenditure through regulation of uncoupling protein 1 expression^14^. Additionally, thyroid hormones influence hepatic lipid metabolism, including cholesterol synthesis and de novo fatty acid synthesis^15^. They also regulate insulin sensitivity and glucose transporter expression, serving as master regulators of whole-body energy homeostasis.

Recent studies have demonstrated that even within the normal thyroid function range, subtle changes in hormone levels are associated with obesity and metabolic abnormalities. The SPECT-China study, which included 8,727 Chinese adults, found that TSH levels within the normal reference range were significantly positively correlated with VAI and LAP (β=0.041 and 0.028, respectively, both P<0.01) ^16^. This suggests that even minor changes in thyroid function may represent potential risk factors for cardiovascular disease.

Weight gain, lipid metabolism disorders, and insulin resistance are commonly observed in patients with hypothyroidism. Research by Pekgor et al. found that VAI levels were significantly higher in both overt and subclinical hypothyroidism patients compared to controls (P<0.01 and P=0.02, respectively), with positive correlations observed between TSH and both BMI and VAI^17^. Recent studies have also demonstrated that abnormal thyroid hormone sensitivity is closely associated with visceral obesity. In 750 euthyroid patients with type 2 diabetes, the peripheral thyroid hormone sensitivity indicator free T3/free T4 ratio was positively correlated with visceral fat area, with moderate and highest tertile groups showing 134% and 98% increased risks of visceral obesity, respectively^18^. The FT3/FT4 ratio reflects peripheral deiodinase activity and tissue thyroid hormone sensitivity, with elevated ratios potentially indicating compensatory responses to metabolic stress.

The association between thyroid function and visceral fat accumulation involves multiple molecular mechanisms: leptin secreted by visceral adipose tissue can induce TRH expression and activate TSH and thyroid hormone synthesis, but leptin resistance in obesity may lead to HPT axis dysfunction^19^; differential expression of deiodinases in various tissues affects local thyroid hormone activity^20^; pro-inflammatory factors produced by visceral fat may interfere with thyroid hormone signal transduction, leading to peripheral thyroid hormone resistance^21^; thyroid hormones also regulate insulin sensitivity through effects on glucose transporter protein expression.

Although previous studies have explored the relationship between thyroid function and obesity, research on the correlation between thyroid function and emerging visceral fat accumulation indices remains relatively limited, particularly systematic analyses based on large population samples. Most existing studies have examined single parameters or simple anthropometric measures without comprehensively evaluating multiple visceral adiposity indices. Different visceral fat accumulation indices may exhibit distinct association patterns with thyroid function parameters, and these associations may demonstrate non-linear characteristics and be modulated by multiple factors.

Based on these considerations, this study utilized the 2007-2012 National Health and Nutrition Examination Survey (NHANES) database to investigate the correlations between visceral fat accumulation indices (METS-VF, LAP, VAI) and thyroid function parameters (TSH, free T3, free T4, total T3, total T4), analyze the linear and non-linear characteristics of these associations, and identify potential modifying factors. We hypothesize that different visceral fat accumulation indices may exhibit specific association patterns with different thyroid function parameters, and that these associations may demonstrate non-linear characteristics modulated by factors such as age, sex, and ethnicity. This research will contribute to a deeper understanding of the mechanisms underlying obesity-related thyroid dysfunction and provide theoretical foundations for clinical early identification and intervention.

## 2 Data and Methods

### 2.1 Data Source

The NHANES data were accessed for research purposes in 27 April, 2025. Data from three survey cycles (2007-2008, 2009-2010, 2011-2012) of the NHANES (https://wwwn.cdc.gov/nchs/nhanes/Default.aspx) were utilized in this study. NHANES is a periodic cross-sectional survey conducted by the Centers for Disease Control and Prevention of the United States, designed to monitor trends in health and nutritional status among the non-institutionalized civilian population of the United States. Through the implementation of a complex, multistage probability sampling design, NHANES provides prevalence estimates for common diseases. The research protocols for NHANES 2007-2010 (Protocol 2005-2006) and 2011-2016 (Protocol 2011-2017) were approved by the NCHS Research Ethics Review Board, and written informed consent was obtained from all participants. The authors had no access to any information that could identify individual participants during or after data collection.

### 2.2 Study Population

This study included three data cycles from the database, namely 2007-2008, 2009-2010, and 2011-2012. From 30,442 subjects, 22,697 participants lacking lipid accumulation indicators were excluded, along with 3,607 lacking relevant thyroid function information and 566 participants under 18 years of age. Subsequently, 1,178 participants with missing data for other covariates were excluded. Ultimately, 2,394 participants were included in this study (Figure 1).

**Figure 1.**
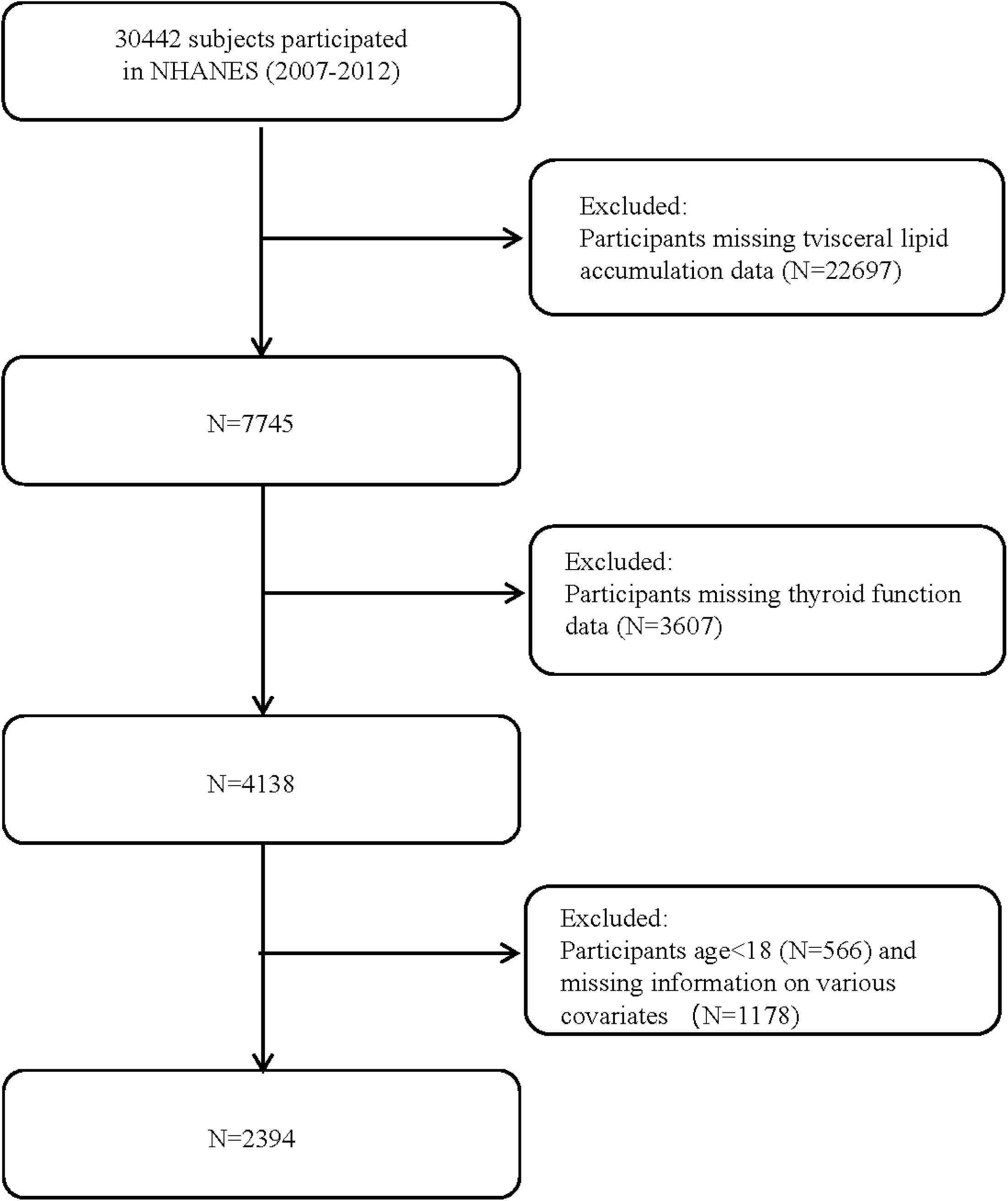
Flow chart of inclusion and exclusion process for NHANES participants from 2007-2012.

### 2.3 Serum Thyroid Measurements

Thyroid examination comprises a series of tests designed to evaluate thyroid activity. This panel includes thyroid-stimulating hormone (TSH), total thyroxine (TT4), free thyroxine (FT4), total triiodothyronine (TT3), free triiodothyronine (FT3), thyroglobulin (Tg), thyroglobulin antibody (TgAb), and thyroid peroxidase antibody (TPOAb). Thyroid blood specimens were processed, stored, and transported to the University of Washington in Seattle, Washington. Detailed specimen collection and processing instructions are discussed in the NHANES Laboratory/Medical Technologists Procedures Manual (LPM).

### 2.4 Lipid Accumulation Indices

METS-VF, VAI, and LAP were considered as exposure variables. Their calculations were determined by the following formulas^11, 12, 22–24^. At the Mobile Examination Center (MEC), professional health technicians carefully measured individuals’ BMI, WC, and height. Data for high-density lipoprotein cholesterol (HDL-C) and triglycerides (TG) were obtained using the Cobas 6000 biochemical analyzer. Data for fasting blood glucose (FBG) were obtained with the Roche/Hitachi Cobas C chemistry analyzer - C311.

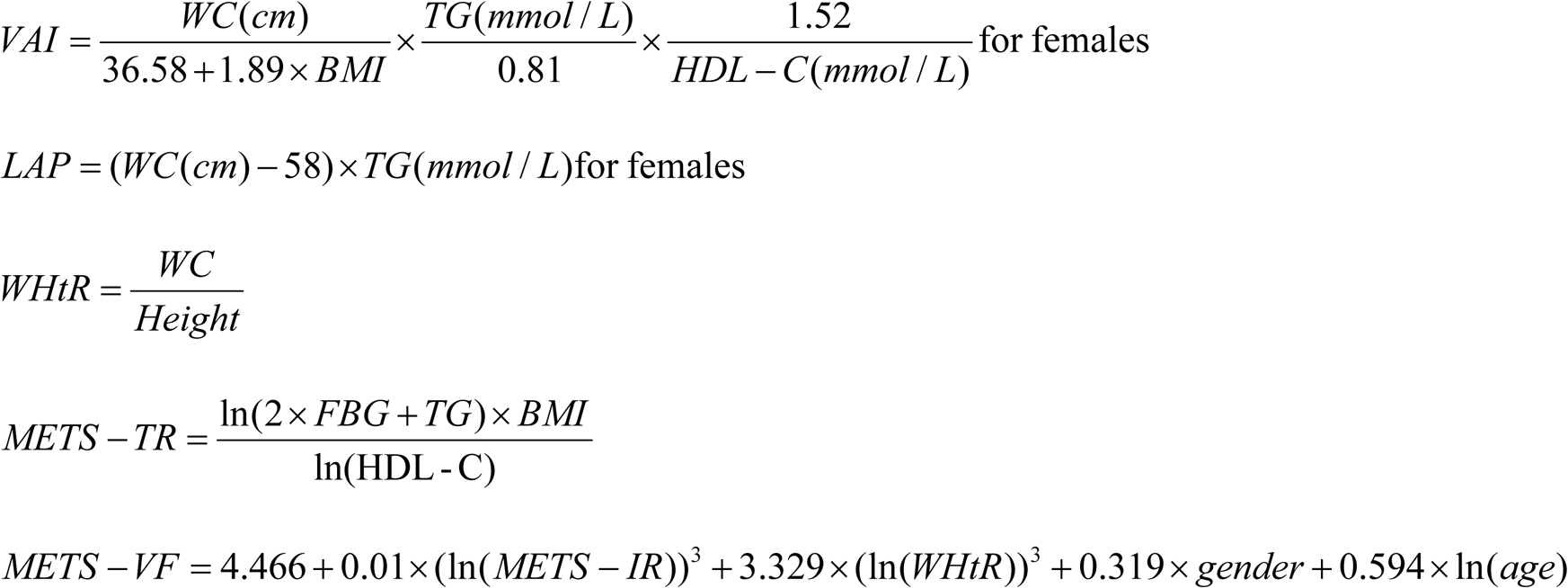

### 2.5 Covariates

Multiple covariates were evaluated for their potential influence based on existing literature. Covariates included age, race/ethnicity, education level, marital status, household income, body mass index (BMI), hypertension, diabetes, smoking status, and alcohol consumption. Race/ethnicity categories were classified as Mexican American, Spanish, Non-Hispanic White, Non-Hispanic Black, and Other Races. Education levels were divided into four groups (elementary school, middle school, high school, and college). Marital status was categorized as married, living with partner, divorced, never married, separated, or widowed. Based on United States government standards, household income was classified as low (≤1.3), medium (>1.3 to 3.5), or high (>3.5) according to the poverty income ratio (PIR). BMI was calculated from examination data. Hypertension and diabetes history were self-reported by participants through questionnaires. Smoking status was categorized as never smokers (<100 cigarettes) or current smokers (≥100 cigarettes)^25^. Alcohol status was defined as at least 12 drinks in a lifetime. Given the importance of iodine for thyroxine synthesis, the fact that iron deficiency impairs thyroid metabolism, and the correlation between thyroid function and renal function, urinary iodine concentration (UIC), creatinine (Cre), and dietary micronutrient intakes of zinc (Zn), selenium (Se), sodium (Na), and potassium (K) were also included in our study.

### 2.6 Statistical Analysis

All statistical analyses were conducted using R software packages. Continuous variables were represented as means (standard deviation, SD), while categorical variables were summarized as frequencies and percentages. For baseline characteristics, comparisons were made using chi-square tests for categorical variables and one-way analysis of variance for continuous variables. Multivariate linear regression models were implemented to analyze potential correlations between lipid accumulation and thyroid function indicators. Model Ⅰ was not adjusted for any covariates, while Model Ⅱ was adjusted for a comprehensive range of factors including age, sex, education level, and marital status. Building upon Model Ⅱ, Model Ⅲ further adjusted for additional confounding factors, including household income, alcohol consumption, smoking, hypertension, diabetes, Zn, Se, Na, K, UIC, and Cre. Subgroup analyses, interaction tests, and restricted cubic spline (RCS) regression were employed to evaluate non-linear associations between lipid accumulation and thyroid function. *P* < 0.05 was considered statistically significant.

## 3 Results

### 3.1 Baseline Participant Characteristics

This study, based on NHANES (2007-2012) data, included 2394 participants who were grouped by LAP (visceral adiposity index) quartiles. The research objective was to analyze differences in demographic, socioeconomic, dietary status, and thyroid function variables among different LAP quartile groups. Table 1 displays the baseline characteristics of these participants. Higher LAP levels were associated with increased age, higher proportion of females, lower education levels, and higher smoking rates. As shown in Table 1, thyroid function indicators with statistically significant differences across LAP quartiles included FT4, FT3/FT4, TSH, TT3, and TT4 (*P* < 0.05). Among covariates, all showed statistical significance (*P* < 0.05) except for Zn, K, Cre, and income level, which did not demonstrate statistical differences (*P* > 0.05).

**Table 1.**
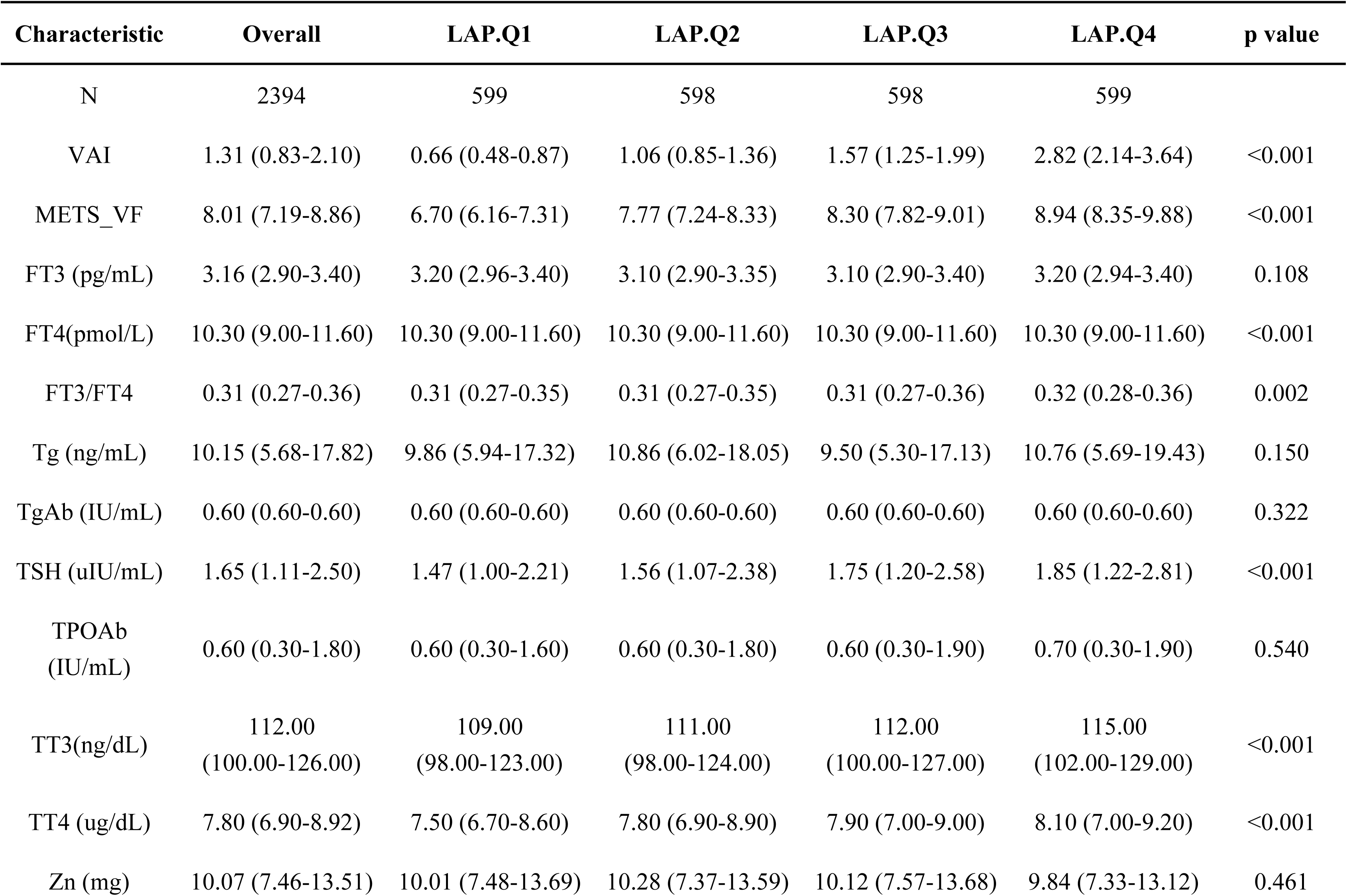
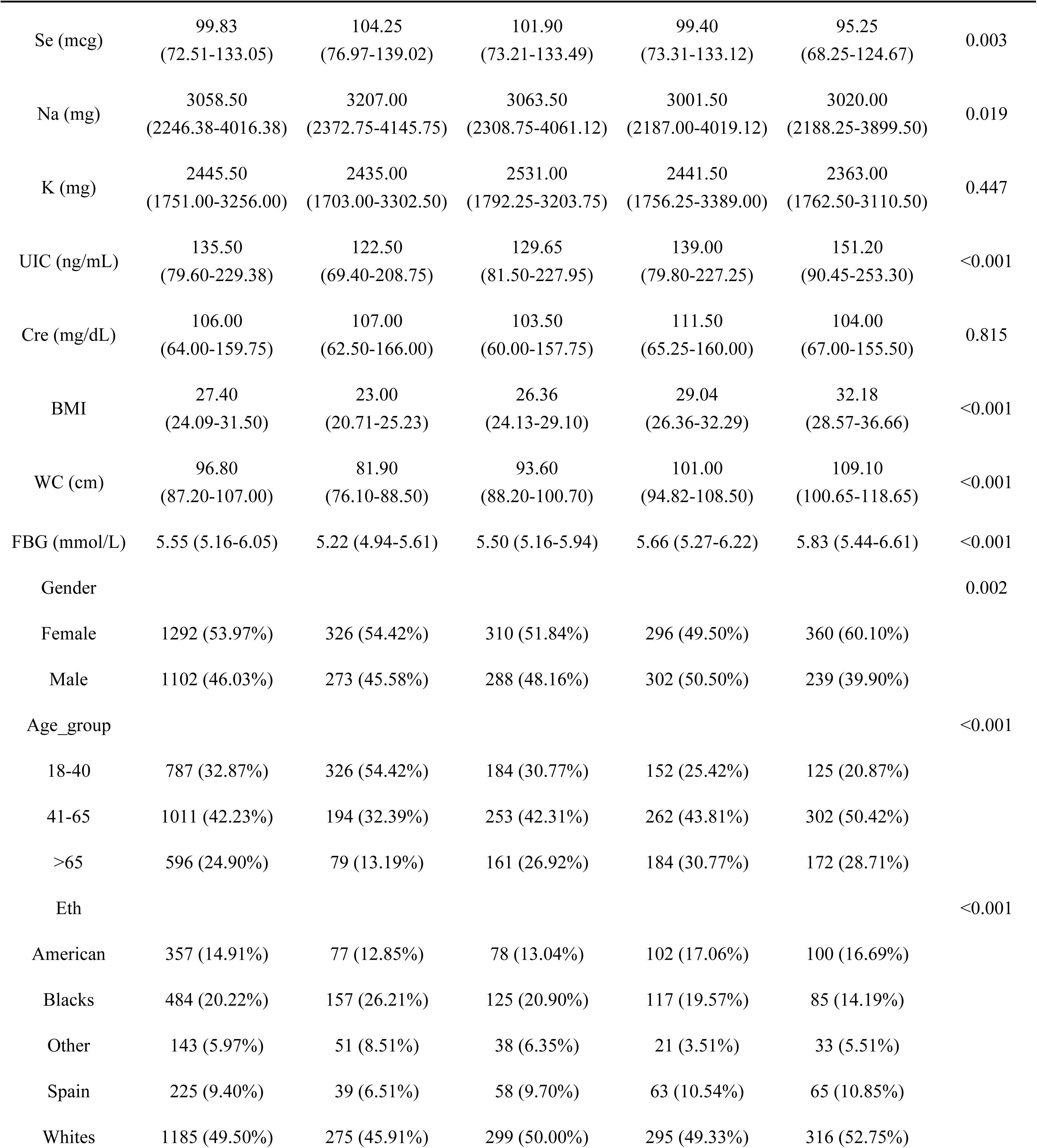
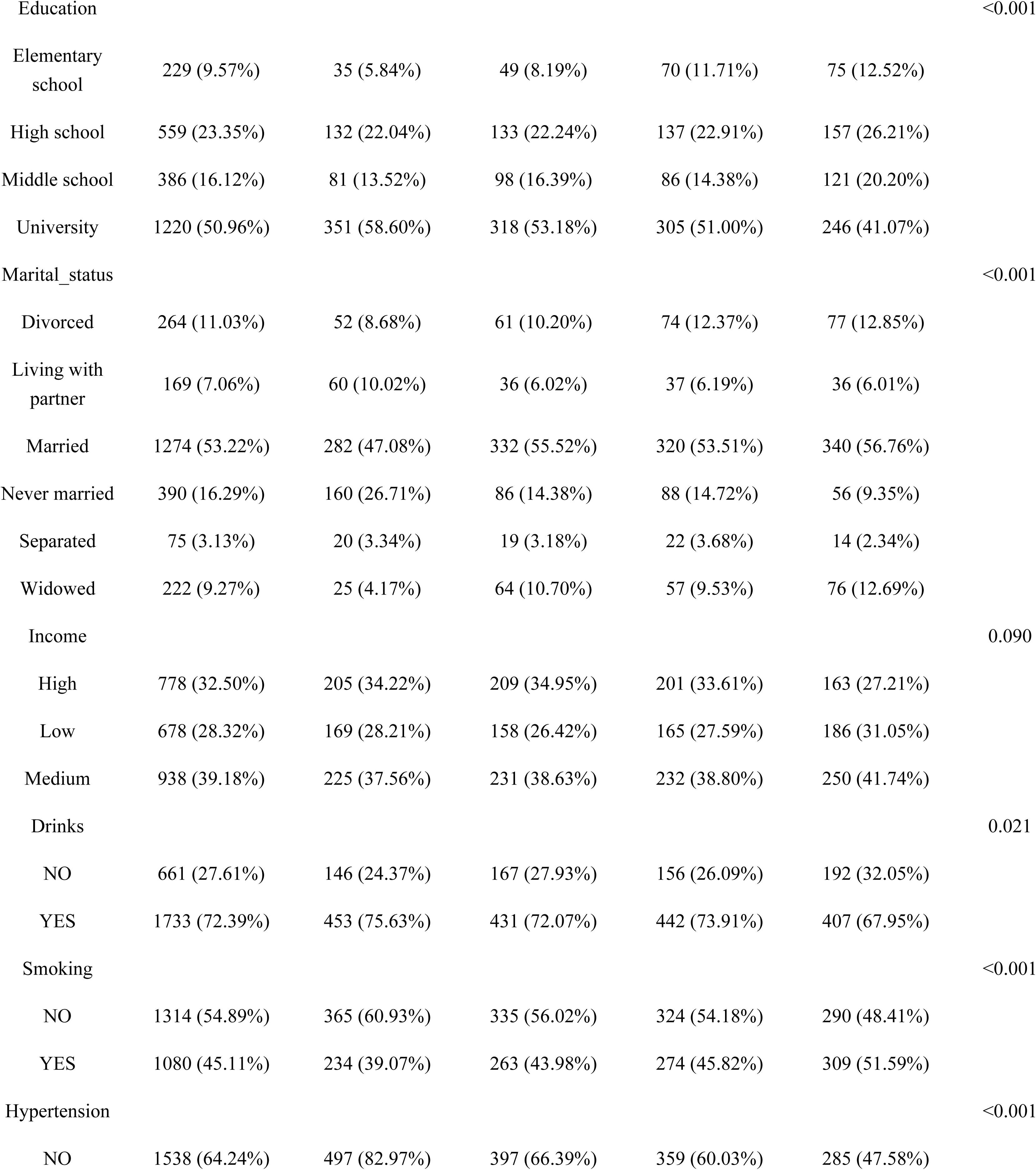
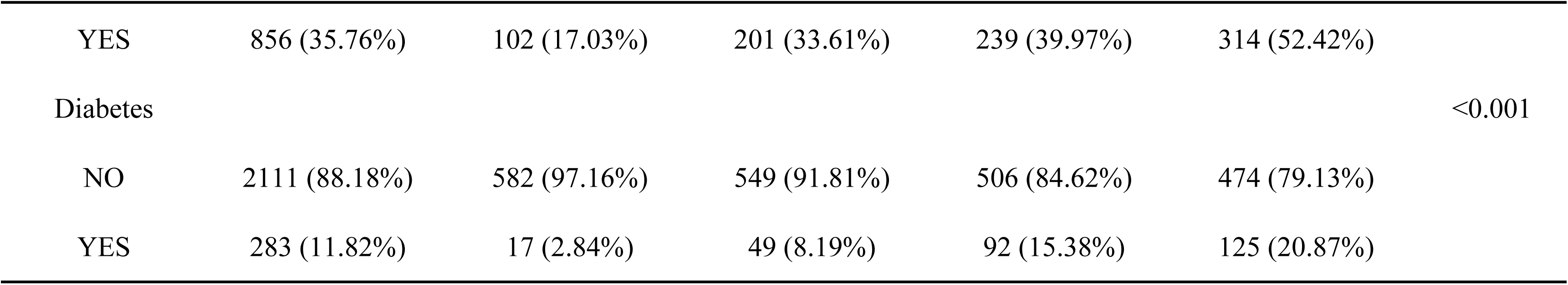
Baseline Participant Characteristics.

### 3.2 Relationship Between Lipid Accumulation and Thyroid Function

To further explore the associations between visceral adiposity indices and thyroid function, we performed multivariate linear regression analyses (Supplementary Tables S1-S3). In the fully adjusted model (Model III), all three indices demonstrated significant positive associations with FT3, FT3/FT4 ratio, TT3, TT4, and TgAb when analyzed as continuous variables (per IQR increase). Notably, both LAP (β = −0.218, P < 0.001) and VAI (β = −0.183, P < 0.001) showed inverse associations with FT4, whereas METS-VF exhibited no significant relationship (β = −0.015, P = 0.746). Associations with TSH were significant for VAI (β = 0.149, P = 0.041) and METS-VF (β = 0.167, P = 0.025), while LAP showed a borderline effect (β = 0.135, P = 0.082). Among the three indices, only METS-VF demonstrated a significant positive association with Tg (β = 2.712, P = 0.031). No significant associations were observed with TPOAb for any index in continuous analysis. Quartile analyses revealed more pronounced associations. Compared with the lowest quartile (Q1), participants in the highest quartile (Q4) showed consistent positive associations with FT3, FT3/FT4 ratio, TT3, TT4, and TgAb across all indices (all P < 0.05). The inverse association with FT4 persisted for LAP (β = −0.660, P < 0.001) and VAI (β = −0.430, P < 0.001) but not METS-VF (P = 0.535). TSH showed significant associations in quartile comparisons for LAP (β = 0.431, P = 0.029) and VAI (β = 0.579, P = 0.003), but the relationship weakened for METS-VF (P = 0.208). Dose-response trend tests confirmed significant linear relationships for FT3, FT3/FT4 ratio, TT3, TT4, and TgAb across all indices (all P for trend<0.05). LAP additionally showed a marginally significant trend for TPOAb (P for trend=0.045), while METS-VF demonstrated a borderline trend for Tg (P for trend=0.058). No significant trends were observed for Tg with LAP (P = 0.512) and VAI (P = 0.782), or for TPOAb with VAI (P = 0.177) and METS-VF (P = 0.522).

### 3.3 RCS Analysis

Restricted cubic spline (RCS) analysis was performed to examine potential non-linear associations between visceral adiposity indices and thyroid function parameters after adjusting for confounders. For LAP (Figure 2), significant non-linear relationships were observed with FT3, TSH, TT3, and TT4 (all P-nonlinear<0.05), while FT4 showed a consistent inverse linear trend (P-overall<0.001, P-nonlinear>0.05). Associations with Tg and TPOAb were non-significant, whereas TgAb demonstrated a weak non-linear pattern.

**Figure 2.**
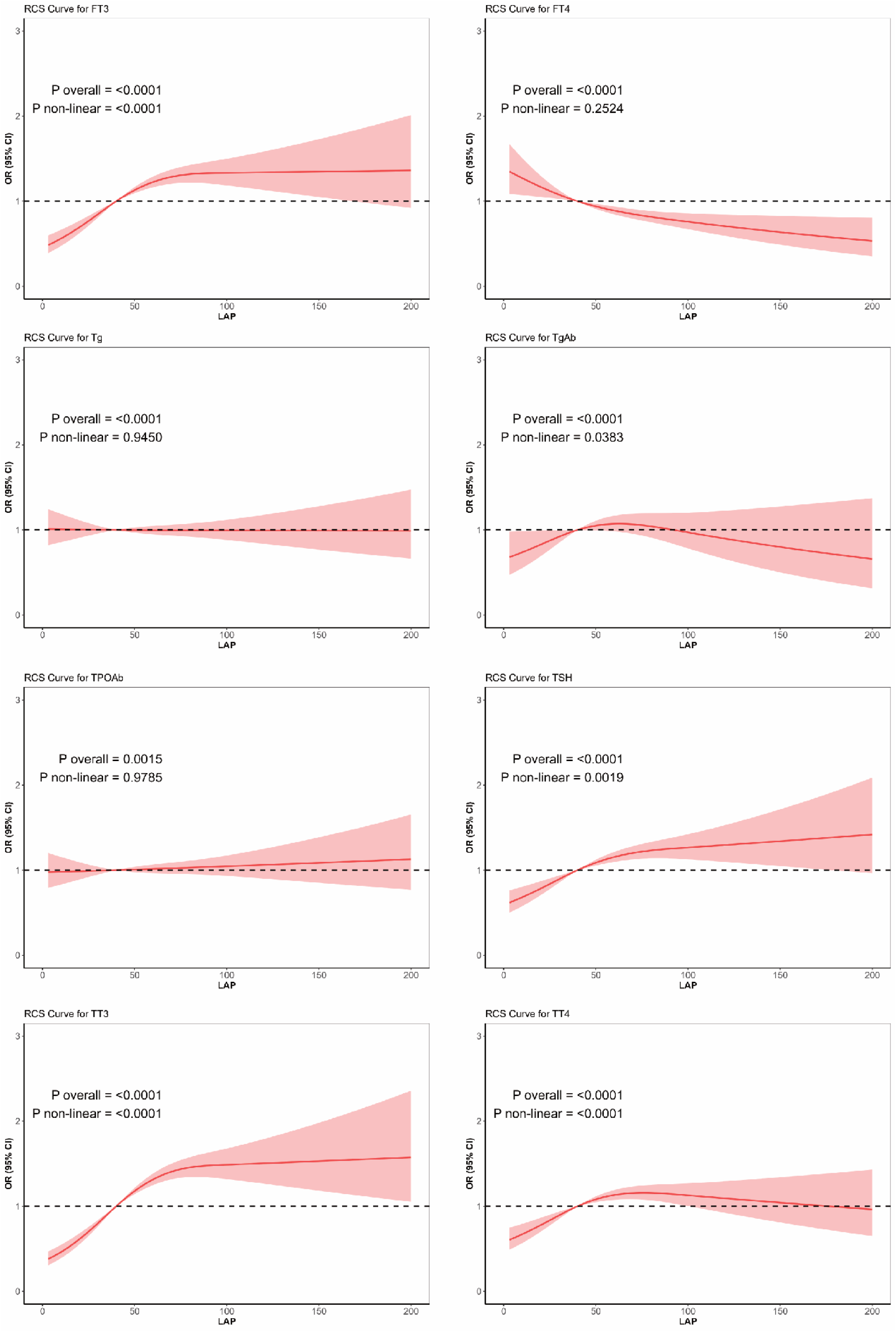
Restricted cubic spline (RCS) regression analysis curves between lipid accumulation product (LAP) and various thyroid function indicators. The RCS curves illustrate the associations between LAP and FT3, FT4, FT3/FT4, TSH, TT3, TT4, Tg, TgAb, and TPOAb. The red solid line represents the OR (and its 95% confidence interval), with the shaded area showing the 95% confidence interval. *P* values are presented for overall association and non-linear trend tests.

VAI exhibited distinct curve patterns across thyroid parameters (Figure 3). FT3, TSH, and TT3 displayed reverse L-shaped curves with rapid initial increases followed by plateaus, while TT4 demonstrated a subtle inverted U-shaped relationship. In contrast, FT4, Tg, TgAb, and TPOAb showed predominantly linear associations with modest effect sizes.

**Figure 3.**
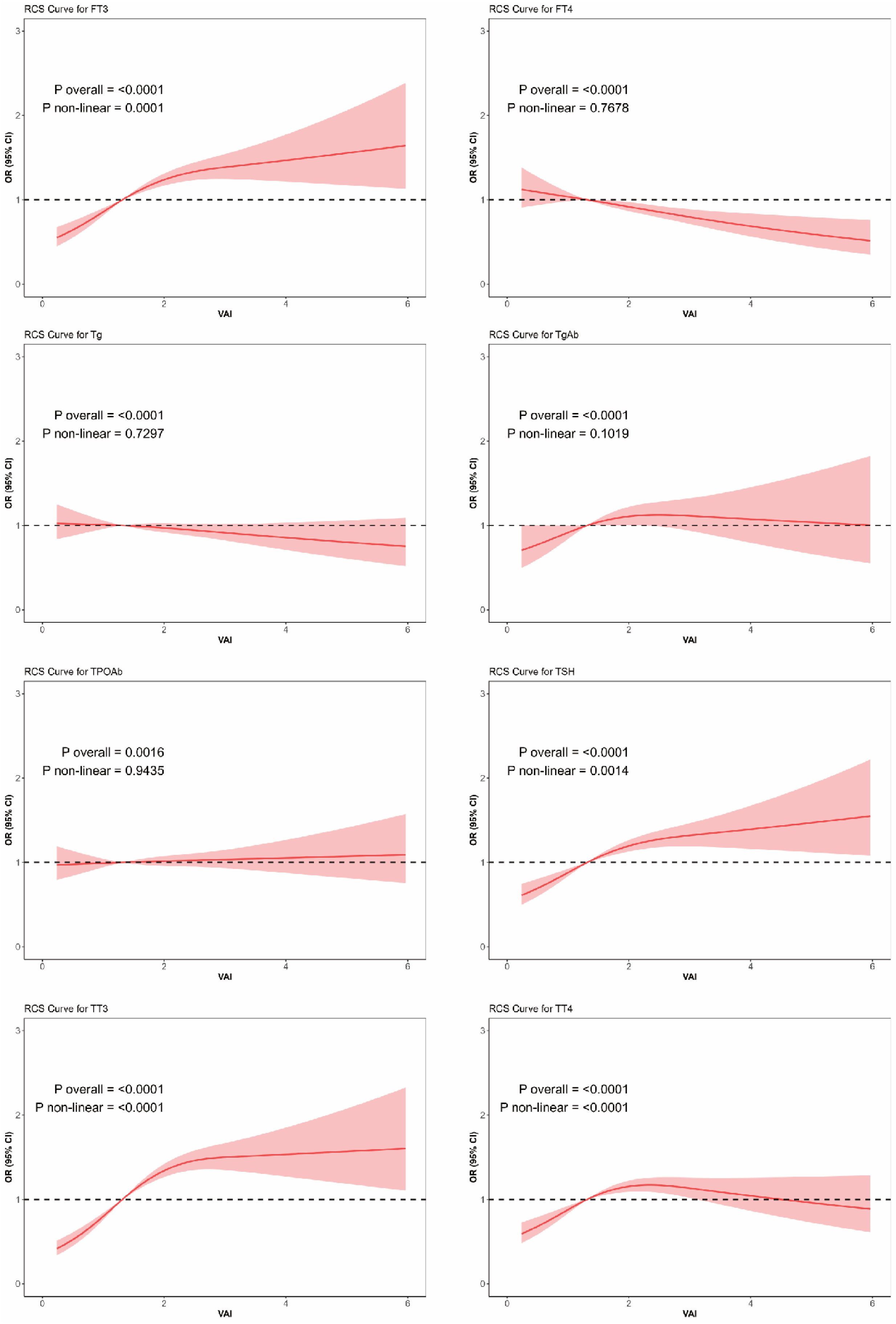
RCS regression analysis curves between visceral adiposity index (VAI) and various thyroid function indicators. The RCS curves illustrate the relationships between VAI and FT3, FT4, FT3/FT4, TSH, TT3, TT4, Tg, TgAb, and TPOAb. The red solid line and shaded area represent the OR and its 95% confidence interval, respectively. *P* values are presented for overall association and non-linear trend tests.

METS-VF revealed heterogeneous association patterns (Figure 4). FT3 and TT3 exhibited pronounced non-linear increases with rising METS-VF, whereas FT4 showed a slight inverted U-shaped curve suggesting a potential optimal range. TSH and TT4 demonstrated linear positive associations. Tg, TgAb, and TPOAb showed weak associations without significant non-linearity.

**Figure 4.**
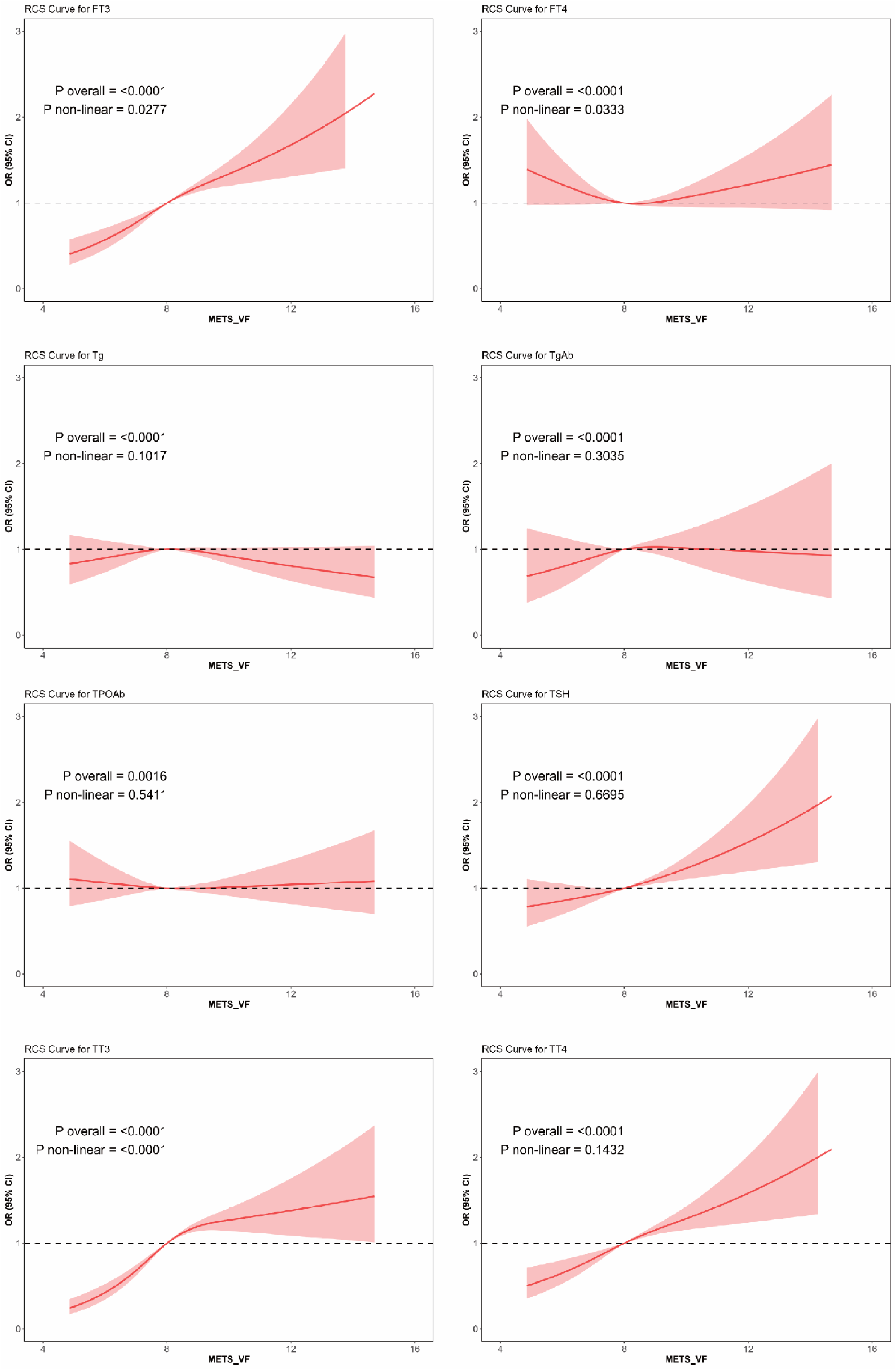
RCS regression analysis curves between metabolic syndrome-related visceral fat (METS-VF) and various thyroid function indicators. The RCS curves reflect the associations between METS-VF and FT3, FT4, FT3/FT4, TSH, TT3, TT4, Tg, TgAb, and TPOAb. The red line and shaded area represent the OR and 95% confidence interval. *P* values are presented for overall and non-linear trend test results.

### 3.4 Subgroup Analysis

Subgroup analyses were performed using multivariate linear regression stratified by age, gender, race, smoking status, alcohol consumption, household income, hypertension, and diabetes to identify potential effect modifiers. Age, race, and smoking consistently modified associations between all three visceral adiposity indices (LAP, VAI, METS-VF) and FT3/FT4 ratio (all interaction P<0.05), with stronger effects observed in younger adults (18-40 years), older adults (>65 years), males, White participants, and high-income groups (Figures 5A, 6A, 7A). For LAP specifically, these associations were also pronounced among alcohol consumers, while VAI showed enhanced effects in individuals with diabetes. In contrast, effect modification patterns for TT3 varied across indices: age emerged as the primary modifier for LAP-TT3 associations (Figure 5B), whereas gender, race, and diabetes modified VAI-TT3 relationships (Figure 6B), and notably, METS-VF showed no significant associations with TT3 across any subgroup examined (Figure 7B). Similarly, TT4 associations demonstrated index-specific modification patterns, with gender, race, and household income modifying LAP-TT4 relationships (Figure 5C), age serving as the key modifier for VAI-TT4 (Figure 6C), and gender and race modifying METS-VF-TT4 associations (Figure 7C). These findings collectively suggest that demographic and lifestyle factors differentially modulate the relationships between visceral adiposity and thyroid function, with consistent modifiers identified for peripheral thyroid hormone ratios but heterogeneous patterns observed for individual hormone levels across different adiposity indices.

**Figure 5.**
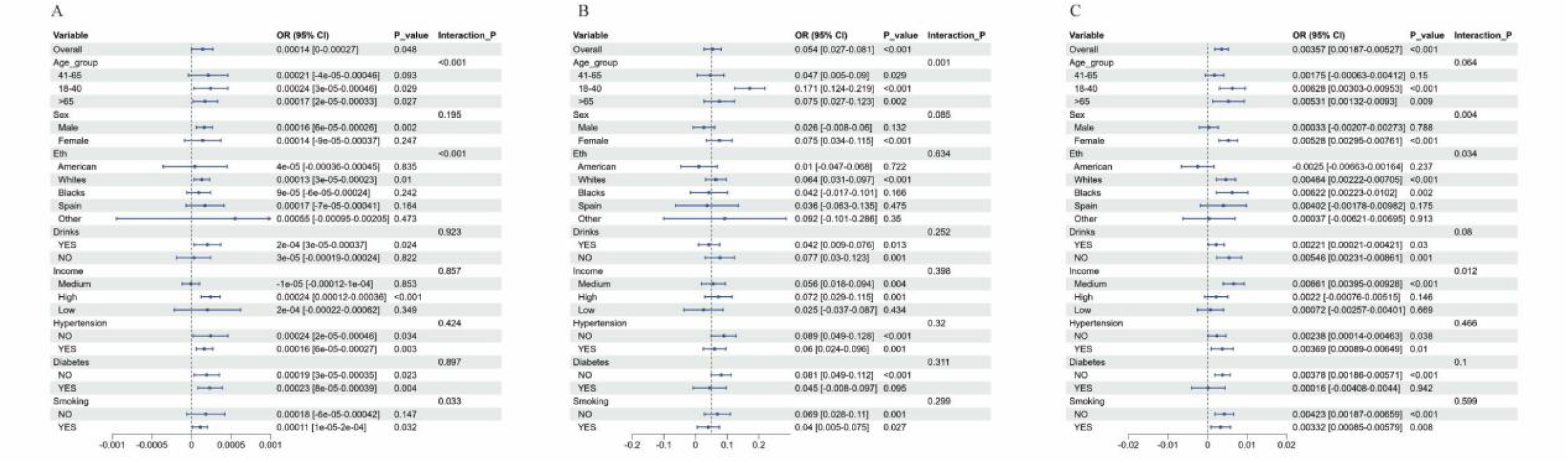
Multivariate linear regression subgroup analysis of associations between LAP and thyroid function indicators across different subgroups. (A) Association between LAP and FT3/FT4 ratio; (**B)** Association between LAP and TT3; (**C)** Association between LAP and TT4.

**Figure 6.**
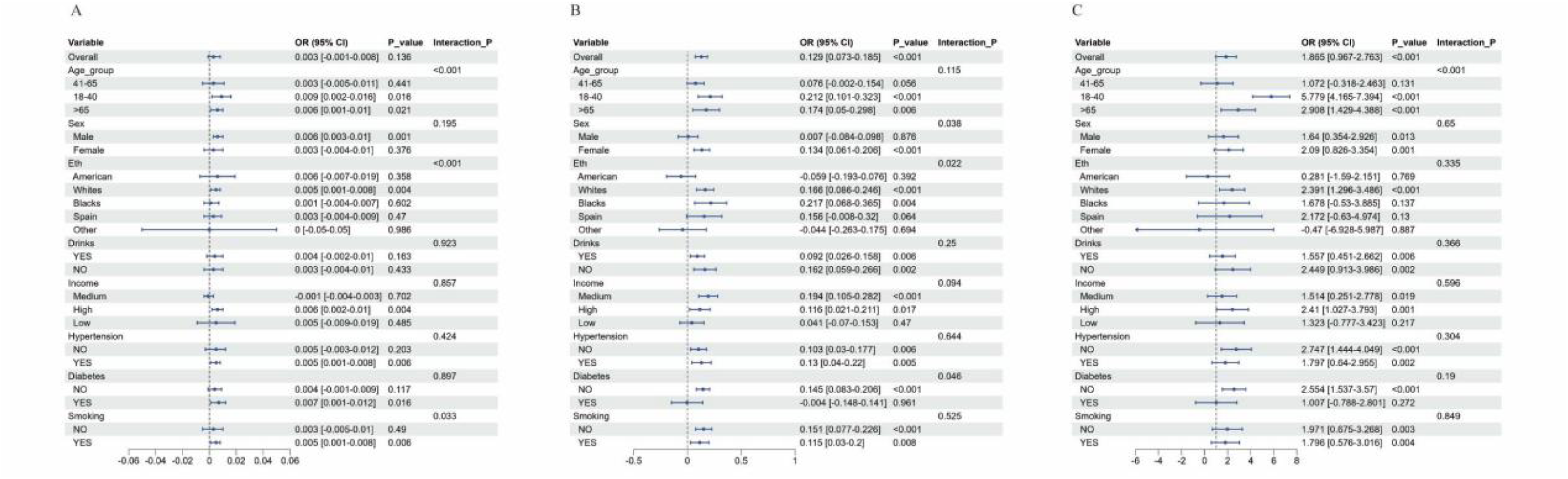
Multivariate linear regression subgroup analysis of associations between VAI and thyroid function indicators across different subgroups. **(A)** Association between VAI and FT3/FT4 ratio; (**B)** Association between VAI and TT3; (**C)** Association between VAI and TT4.

**Figure 7.**
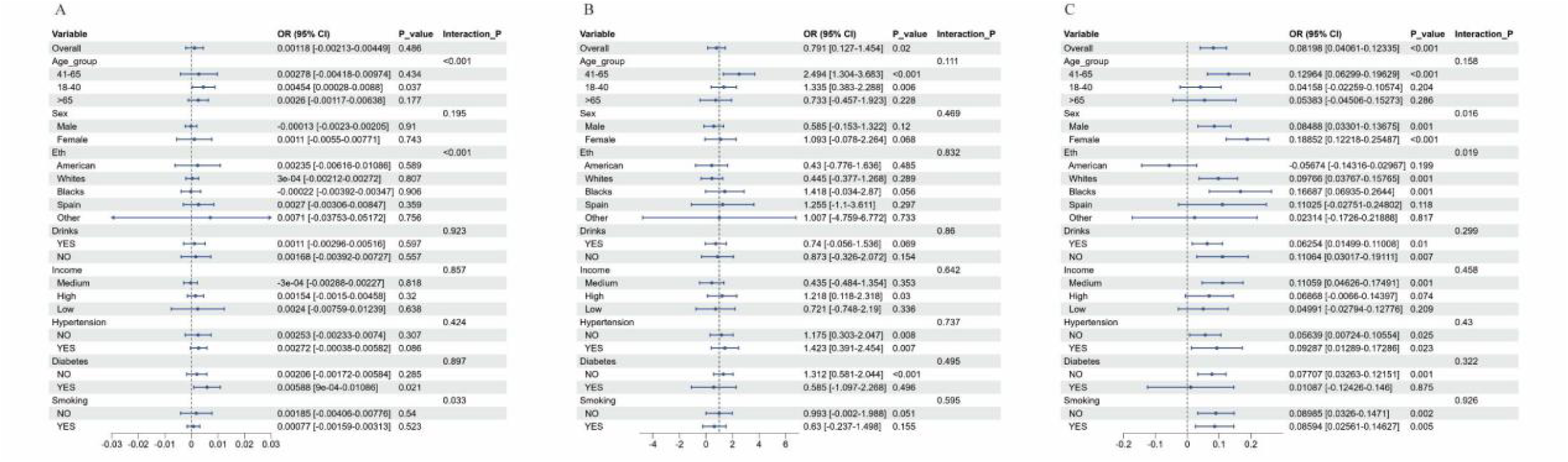
Multivariate linear regression subgroup analysis of associations between METS-VF and thyroid function indicators across different subgroups. **(A)** Association between METS-VF and FT3/FT4 ratio; (**B)** Association between METS-VF and TT3; (**C)** Association between METS-VF and TT4.

## 4 Discussion

This study, based on data from the NHANES 2007-2012, systematically explored the associations between visceral adiposity indices and thyroid function parameters. Through multiple statistical analytical methods, we identified significant associations between visceral adiposity and thyroid function, which exhibited complex nonlinear characteristics and were modulated by factors such as age, race, and lifestyle habits. The core findings of this study encompass three main aspects. After adjusting for potential confounding factors, three visceral adiposity indices (METS-VF, LAP, and VAI) demonstrated positive correlations with FT3, FT3/FT4 ratio, TSH, TT3, and TT4, while showing negative correlations with FT4. These findings are consistent with the retrospective cohort study conducted by Mahdavi et al., which identified a positive correlation between obesity and hypothyroidism risk^26^. Notably, another systematic review and meta-analysis based on prospective studies revealed that obese individuals at baseline had significantly increased risks of developing overt hypothyroidism (RR: 3.10, 95% CI 1.56–4.64, I² = 0%) and subclinical hypothyroidism (RR 1.50, 95% CI 1.05–1.94, I² = 0%)^27^. These quantitative data provide robust epidemiological support for the associations we observed between visceral adiposity indices and thyroid function parameters. Furthermore, causal inference analyses employing Mendelian randomization methodology demonstrated a positive causal association between obesity and hypothyroidism^28^, providing causal evidence to support the correlations identified in previous observational studies. This multilayered evidence chain supports that visceral fat accumulation is indeed an independent risk factor for thyroid dysfunction.

Compared to the traditional BMI metric, the visceral adiposity indices employed in this study—LAP, VAI, and METS-VF—can more precisely reflect metabolic abnormalities. A large-scale cross-sectional study in the Chinese population found that even within the reference range of TSH, higher TSH levels were significantly associated with VAI and LAP^16^, supporting the rationale for using visceral adiposity indices rather than simple BMI in this study. Another investigation further confirmed that increased thyroid hormone sensitivity is closely associated with visceral obesity. That study, using multivariable regression analysis, demonstrated that the prevalence of visceral obesity in the middle and high tertiles of FT3/FT4 ratio was 134% and 98% higher, respectively, compared to the lowest tertile^18^. This finding is highly consistent with the positive correlation between FT3/FT4 ratio and visceral adiposity indices observed in our study. Notably, multiple studies on childhood and adolescent obesity have likewise confirmed similar association patterns. Research has revealed elevated TSH, T3, and FT3 levels with normal or decreased T4 in obese children^29, 30^. These early-onset metabolic alterations suggest that the impact of visceral fat accumulation on thyroid function may begin in childhood and persist into adulthood. Through restricted cubic spline (RCS) analysis, this study revealed that these associations predominantly exhibited nonlinear patterns. Significant "inverted L-shaped" curves were observed between LAP and FT3, TSH, TT3, and TT4. VAI and METS-VF demonstrated similar nonlinear patterns. This finding aligns with previous reports identifying nonlinear associations between body roundness index (BRI) and thyroid hormones with threshold effects (BRI=7.21)^31^. These nonlinear characteristics and threshold effects provide quantitative evidence for clinical assessment and suggest that the body may undergo dynamic compensatory-decompensatory processes at different visceral fat levels.

The "inverted L-shaped" nonlinear curve observed in this study may reflect the dynamic compensatory-decompensatory processes of the body at different visceral fat levels. During stages of lower visceral fat content, the body may maintain relatively stable thyroid function through multiple compensatory mechanisms: regulation of thyroid hormone receptor expression, adjustment of deiodinase activity, and feedback regulation of the hypothalamic-pituitary-thyroid axis^32^. However, when visceral fat exceeds a specific threshold, the association between visceral adiposity and thyroid dysfunction may be mediated through multiple mechanisms. First, visceral fat, as an active endocrine organ, secretes various inflammatory cytokines such as TNF-α, IL-6, and IL-1β, which subsequently affect the function of the hypothalamic-pituitary-thyroid axis. These inflammatory cytokines may lead to elevated TSH and decreased FT4 by suppressing the synthesis and release of TRH and TSH, interfering with thyroid hormone receptor function, or affecting peripheral conversion of thyroid hormones^33^. Second, previous studies have demonstrated that visceral fat accumulation is closely associated with insulin resistance, and insulin resistance may influence thyroid hormone metabolism and peripheral conversion^34^. Insulin resistance can promote the conversion of T4 to T3, increase type 1 deiodinase (D1) activity, while modulating type 3 deiodinase (D3) activity to affect T3 metabolism^34^, which may explain the positive correlation between FT3/FT4 ratio and visceral adiposity indices observed in our study. Adipokines produced by visceral fat, such as leptin and adiponectin, may directly influence thyroid function. Leptin may enhance central nervous system regulation of the thyroid axis by stimulating the synthesis and release of TRH and TSH^35^, while decreased adiponectin levels may be associated with hypothyroidism^36^. The dysregulation of these adipokines may represent an important pathway through which visceral adiposity affects thyroid function.

The findings of this study have important clinical implications. First, for patients with visceral obesity, routine screening of thyroid function should be considered, particularly TSH, FT4, and TT3 levels, to enable early identification of potential thyroid dysfunction. Second, when visceral fat accumulation exceeds a certain threshold, the risk of thyroid dysfunction may increase significantly. Clinicians may need to pay particular attention to patients whose visceral adiposity indices fall in the highest quartile, as their risk of thyroid dysfunction may increase nonlinearly. Third, subgroup analysis results indicate that age, race, and lifestyle habits play modulatory roles in the association between visceral adiposity and thyroid function, providing a foundation for individualized risk assessment. For instance, we found that associations between visceral adiposity and thyroid dysfunction were more pronounced in the 18-40 years and >65 years age groups, White race, and smokers, suggesting these populations may require more aggressive thyroid function screening and intervention strategies. Finally, the study suggests that reducing visceral fat may be an effective approach to improving obesity-related thyroid dysfunction. Clinicians may need to emphasize strategies targeting visceral fat reduction in treatment plans, such as lifestyle interventions, pharmacotherapy, or surgical procedures, which may simultaneously improve metabolic health and thyroid function.

However, this study has several limitations. First, as a cross-sectional study design, we cannot establish causal relationships between visceral adiposity and thyroid dysfunction. Second, although we adjusted for multiple potential confounding factors, unmeasured confounders may still influence the study results. Third, the visceral adiposity indices in this study (LAP, VAI, and METS-VF) were indirectly calculated based on anthropometric and biochemical indicators, rather than directly measured using gold-standard methods for visceral fat content such as CT or MRI. This may introduce measurement error and affect estimates of association strength. Finally, due to the geographic limitations of NHANES data, the study results are primarily applicable to the US population, and the generalizability to other racial and regional populations requires further validation. Particularly considering that different races may exhibit variations in thyroid function and obesity characteristics, the study results should be interpreted cautiously when applied to other populations.

Based on the findings and limitations of this study, we propose several future research directions. First, prospective cohort studies are needed to track dynamic changes in visceral adiposity and thyroid function, clarifying the causal relationships and temporal characteristics between them. Second, combining advanced imaging technologies (such as CT and MRI) to directly measure visceral fat content and conducting association analyses with thyroid function would overcome the limitations of indirect calculation indices, while integrating molecular biology techniques to explore the molecular mechanisms underlying their associations, such as interactions among adipokines, inflammatory cytokines, and the thyroid axis. Third, the generalizability of this study’s findings should be validated in different racial and regional populations, exploring race-specific association patterns. Particularly in Asian populations, where body fat distribution characteristics differ from Western populations, the associations between visceral adiposity and thyroid function may exhibit unique features. Finally, we recommend developing thyroid dysfunction risk prediction models that integrate visceral adiposity indices and account for nonlinear association characteristics and heterogeneity across different population subgroups, to enhance prediction accuracy and individualization.

## 5 Conclusion

In summary, this study, based on NHANES 2007-2012 data, systematically revealed significant associations between visceral adiposity and thyroid function. After adjusting for potential confounding factors, LAP, VAI, and METS-VF demonstrated positive correlations with FT3, FT3/FT4 ratio, TSH, TT3, and TT4, and negative correlations with FT4, with these associations predominantly exhibiting nonlinear patterns. Age, race, and smoking play important roles in modulating these associations. These findings provide new perspectives for understanding the complex relationship between obesity and thyroid function, emphasizing the importance of considering visceral fat distribution when assessing thyroid health. Future research should focus on elucidating causal relationships and molecular mechanisms between them, as well as developing individualized thyroid dysfunction risk assessment tools based on visceral adiposity indices, to provide more precise guidance for clinical practice.

## Author Contributions

KC and WP contributed equally to this work. KC: conceptualization, NHANES data extraction and cleaning, statistical analysis, manuscript writing, and visualization. WP: data processing, statistical modeling, result interpretation, and manuscript revision. ZY: study design, methodology guidance, data interpretation, critical revision, and final approval. All authors read and approved the final manuscript.

## Acknowledgments

We would like to express our sincere gratitude to all the participants for their contributions to this study.

## Availability of data

All data used in this study are publicly available. All data used in this study are publicly available from the National Health and Nutrition Examination Survey (NHANES) database. The datasets analyzed during the current study can be accessed through the NHANES website (https://wwwn.cdc.gov/nchs/nhanes/Default.aspx). Specifically, data from three survey cycles (2007-2008, 2009-2010, and 2011-2012) were utilized, including demographic information, anthropometric measurements, laboratory examination results (thyroid function parameters, lipid profiles, glucose metabolism indicators, and micronutrient levels), and questionnaire data. All NHANES data are de-identified and publicly accessible, with detailed documentation available on the NHANES website regarding data collection methods, laboratory procedures, and quality control measures.

## Consent to Publish declaration

Not applicable.

## Ethics and Consent to Participate declaration

Not applicable.

## Clinical trial number

Not applicable.

---

**Supplementary Table S1.** The association between LAP with thyroid function.

| FT3 | Model I | P | Model II | P | Model III | P |
| --- | --- | --- | --- | --- | --- | --- |
| LAP per IQR | -0.008 (-0.039, 0.02) | 0.61 | 0.032 (0.001, 0.06) | 0.042 | 0.039 (0.007, 0.07) | 0.016 |
| Quartiles of LAP |  |  |  |  |  |  |
| Q1 | Ref |  | Ref |  | Ref |  |
| Q2 | -0.08 (-0.16, -0.01) | 0.027 | 0.004 (-0.072, 0.081) | 0.909 | 0.007 (-0.069, 0.084) | 0.856 |
| Q3 | -0.038 (-0.115, 0.04) | 0.343 | 0.068 (-0.009, 0.145) | 0.085 | 0.079 (0.001, 0.156) | 0.047 |
| Q4 | -0.05 (-0.131, 0.025) | 0.181 | 0.076 (-0.002, 0.155) | 0.056 | 0.091 (0.011, 0.171) | 0.026 |
| P for trend | 0.484 |  | 0.027 |  | 0.01 |  |
| FT4 | Model I | P | Model II | P | Model III | P |
| LAP per IQR | -0.155 (-0.247, -0.06) | 0.001 | -0.197 (-0.291, -0.103) | <0.001 | -0.218 (-0.314, -0.122) | <0.001 |
| Quartiles of LAP |  |  |  |  |  |  |
| Q1 | Ref |  | Ref |  | Ref |  |
| Q2 | -0.213 (-0.44, 0.018) | 0.071 | -0.338 (-0.572, -0.10) | 0.005 | -0.337 (-0.57, -0.103) | 0.005 |
| Q3 | -0.195 (-0.42, 0.036) | 0.099 | -0.34 (-0.57, -0.104) | 0.005 | -0.365 (-0.602, -0.13) | 0.003 |
| Q4 | -0.458 (-0.69, -0.22) | <0.001 | -0.624 (-0.864, -0.38) | <0.001 | -0.66 (-0.90, -0.416) | <0.001 |
| P for trend | <0.001 |  | <0.001 |  | <0.001 |  |
| FT3/FT4 | Model I | P | Model II | P | Model III | P |
| LAP per IQR | 0.006 (0, 0.012) | 0.048 | 0.01 (0.004, 0.016) | 0.001 | 0.012 (0.006, 0.019) | <0.001 |
| Quartiles of LAP |  |  |  |  |  |  |
| Q1 | Ref |  | Ref |  | Ref |  |
| Q2 | -0.002 (-0.017, 0.013) | 0.749 | 0.009 (-0.006, 0.024) | 0.258 | 0.009 (-0.006, 0.025) | 0.226 |
| Q3 | 0.01 (-0.005, 0.025) | 0.178 | 0.023 (0.008, 0.038) | 0.003 | 0.025 (0.01, 0.041) | 0.001 |
| Q4 | 0.011 (-0.004, 0.026) | 0.147 | 0.026 (0.011, 0.042) | 0.001 | 0.03 (0.014, 0.046) | <0.001 |
| P for trend | 0.066 |  | 0.001 |  | <0.001 |  |
| Tg | Model I | P | Model II | P | Model III | P |
| LAP per IQR | 2.112 (-0.319, 4.544) | 0.089 | 1.33 (-1.168, 3.828) | 0.297 | 1.484 (-1.093, 4.061) | 0.259 |
| Quartiles of LAP |  |  |  |  |  |  |
| Q1 | Ref |  | Ref |  | Ref |  |
| Q2 | 3.952 (-2.171, 10.07) | 0.206 | 2.933 (-3.312, 9.179) | 0.357 | 3.15 (-3.115, 9.415) | 0.325 |
| Q3 | 1.304 (-4.82, 7.427) | 0.676 | -0.191 (-6.499, 6.117) | 0.953 | -0.142 (-6.505, 6.221) | 0.965 |
| Q4 | 5.168 (-0.953, 11.28) | 0.098 | 2.892 (-3.502, 9.286) | 0.375 | 3.167 (-3.395, 9.73) | 0.344 |
| *P* for trend | 0.173 |  | 0.544 |  | 0.512 |  |
| **TgAb** | **Model I** | ***P*** | **Model II** | ***P*** | **Model III** | ***P*** |
| LAP per IQR | 5.668 (0.986, 10.349) | 0.018 | 4.344 (-0.454, 9.141) | 0.076 | 6.263 (1.319, 11.208) | 0.013 |
| Quartiles of LAP |  |  |  |  |  |  |
| Q1 | Ref |  | Ref |  | Ref |  |
| Q2 | -1.695 (-13.46, 10.07) | 0.778 | -4.659 (-16.632, 7.313) | 0.446 | -3.644 (-15.64, 8.352) | 0.552 |
| Q3 | 6.381 (-5.38, 18.149) | 0.288 | 2.711 (-9.382, 14.804) | 0.66 | 5.338 (-6.846, 17.523) | 0.391 |
| Q4 | 19.597 (7.834, 31.36) | 0.001 | 15.681 (3.424, 27.938) | 0.012 | 20.368 (7.802, 32.934) | 0.002 |
| *P* for trend | <0.001 |  | 0.002 |  | <0.001 |  |
| **TSH** | **Model I** | ***P*** | **Model II** | ***P*** | **Model III** | ***P*** |
| LAP per IQR | 0.186 (0.043, 0.329) | 0.011 | 0.124 (-0.023, 0.271) | 0.098 | 0.135 (-0.017, 0.287) | 0.082 |
| Quartiles of LAP |  |  |  |  |  |  |
| Q1 | Ref |  | Ref |  | Ref |  |
| Q2 | -0.004 (-0.364, 0.357) | 0.983 | -0.115 (-0.482, 0.253) | 0.541 | -0.102 (-0.471, 0.267) | 0.587 |
| Q3 | 0.409 (0.048, 0.769) | 0.026 | 0.272 (-0.099, 0.643) | 0.151 | 0.284 (-0.09, 0.659) | 0.137 |
| Q4 | 0.567 (0.206, 0.927) | 0.002 | 0.386 (0.01, 0.762) | 0.044 | 0.431 (0.045, 0.818) | 0.029 |
| *P* for trend | <0.001 |  | 0.009 |  | 0.006 |  |
| **TPOAb** | **Model I** | ***P*** | **Model II** | ***P*** | **Model III** | ***P*** |
| LAP per IQR | 3.227 (-1.548, 8.002) | 0.185 | 1.997 (-2.891, 6.885) | 0.423 | 2.816 (-2.211, 7.844) | 0.272 |
| Quartiles of LAP |  |  |  |  |  |  |
| Q1 | Ref |  | Ref |  | Ref |  |
| Q2 | 2.275 (-9.746, 14.29) | 0.711 | 1.26 (-10.956, 13.475) | 0.84 | 1.859 (-10.359, 14.07) | 0.766 |
| Q3 | 3.872 (-8.148, 15.89) | 0.528 | 3.683 (-8.656, 16.022) | 0.559 | 4.593 (-7.816, 17.003) | 0.468 |
| Q4 | 12.324 (0.308, 24.34) | 0.045 | 9.587 (-2.919, 22.093) | 0.133 | 12.126 (-0.672, 24.92) | 0.063 |
| *P* for trend | 0.034 |  | 0.101 |  | 0.045 |  |
| **TT3** | **Model I** | ***P*** | **Model II** | ***P*** | **Model III** | ***P*** |
| LAP per IQR | 2.357 (1.163, 3.552) | <0.001 | 3.906 (2.727, 5.084) | <0.001 | 4.343 (3.136, 5.55) | <0.001 |
| Quartiles of LAP |  |  |  |  |  |  |
| Q1 | Ref |  | Ref |  | Ref |  |
| Q2 | 0.936 (-2.07, 3.943) | 0.542 | 5.226 (2.294, 8.157) | <0.001 | 5.434 (2.514, 8.353) | <0.001 |
| Q3 | 4.16 (1.154, 7.167) | 0.007 | 9.392 (6.431, 12.353) | <0.001 | 10.113 (7.148, 13.078) | <0.001 |
| Q4 | 5.876 (2.871, 8.882) | <0.001 | 11.483 (8.481, 14.484) | <0.001 | 12.425 (9.367, 15.483) | <0.001 |
| *P* for trend | <0.001 |  | <0.001 |  | <0.001 |  |
| TT4 | **Model I** | ***P*** | **Model II** | ***P*** | **Model III** | ***P*** |
| LAP per IQR | 0.113 (0.033, 0.192) | 0.005 | 0.156 (0.082, 0.231) | <0.001 | 0.129 (0.054, 0.205) | 0.001 |
| Quartiles of LAP |  |  |  |  |  |  |
| Q1 | Ref |  | Ref |  | Ref |  |
| Q2 | 0.106 (0.028, 0.183) | 0.008 | 0.449 (0.261, 0.636) | <0.001 | 0.394 (0.2, 0.587) | <0.001 |
| Q3 | 0.237 (0.049, 0.424) | 0.014 | 0.236 (0.047, 0.425) | 0.014 | 0.228 (0.04, 0.416) | 0.017 |
| Q4 | 0.315 (0.127, 0.502) | 0.001 | 0.323 (0.132, 0.514) | 0.001 | 0.298 (0.107, 0.489) | 0.002 |
| *P* for trend | 0.001 |  | <0.001 |  | <0.001 |  |

Model I was crude model. Model II was adjusted for age, sex, edu, Marital status. Model III was adjusted for age, sex, edu, Marital status, income, drinks, Zn, Se, Na, k, smoking, hypertension, diabetes ,UIC, Cre.

**Supplementary Table S2.** The association between VAI with thyroid function.

| **FT3** | **Model I** | **P** | **Model II** | **P** | **Model III** | **P** |
| --- | --- | --- | --- | --- | --- | --- |
| VAI per IQR | -0.001 (-0.031, 0.028) | 0.936 | 0.036 (0.007, 0.065) | 0.015 | 0.040 (0.01, 0.069) | 0.009 |
| Quartiles of VAI |  |  |  |  |  |  |
| Q1 | Ref |  | Ref |  | Ref |  |
| Q2 | 0.073 (-0.005, 0.15) | 0.067 | 0.13(0.055, 0.205) | 0.001 | 0.130 (0.054, 0.205) | 0.001 |
| Q3 | 0.007 (-0.071, 0.084) | 0.866 | 0.077(0.002, 0.153) | 0.046 | 0.080 (0.004, 0.156) | 0.039 |
| Q4 | 0.024 (-0.054, 0.102) | 0.543 | 0.132 (0.055, 0.21) | 0.001 | 0.139 (0.060, 0.217) | 0.001 |
| P for trend | 0.948 |  | 0.014 |  | 0.009 |  |
| **FT4** | **Model I** | **P** | **Model II** | **P** | **Model III** | **P** |
| VAI per IQR | -0.130 (-0.217, -0.042) | 0.004 | -0.162 (-0.252, -0.073) | <0.001 | -0.183 (-0.273, -0.092) | <0.001 |
| Quartiles of VAI |  |  |  |  |  |  |
| Q1 | Ref |  | Ref |  | Ref |  |
| Q2 | 0.003 (-0.228, 0.235) | 0.977 | -0.061 (-0.293, 0.17) | 0.603 | -0.063 (-0.294, 0.168) | 0.594 |
| Q3 | -0.017 (-0.249, 0.215) | 0.886 | -0.090 (-0.322, 0.143) | 0.450 | -0.109 (-0.342, 0.123) | 0.356 |
| Q4 | -0.268 (-0.5, -0.037) | 0.023 | -0.372 (-0.61, -0.135) | 0.002 | -0.430 (-0.670, -0.190) | <0.001 |
| P for trend | 0.011 |  | 0.011 |  | <0.001 |  |
| **FT3/FT4** | **Model I** | **P** | **Model II** | **P** | **Model III** | **P** |
| VAI per IQR | 0.004 (-0.001, 0.010) | 0.136 | 0.008 (0.002, 0.014) | 0.007 | 0.009 (0.003, 0.015) | 0.002 |
| Quartiles of VAI |  |  |  |  |  |  |
| Q1 | Ref |  | Ref |  | Ref |  |
| Q2 | 0.014 (-0.001, 0.029) | 0.066 | 0.021 (0.006, 0.036) | 0.007 | 0.021 (0.006, 0.036) | 0.007 |
| Q3 | 0.001 (-0.014, 0.016) | 0.871 | 0.009 (-0.006, 0.024) | 0.239 | 0.010 (-0.005, 0.025) | 0.194 |
| Q4 | 0.013 (-0.002, 0.028) | 0.082 | 0.025 (0.009, 0.04) | 0.002 | 0.027 (0.012, 0.043) | 0.001 |
| P for trend | 0.236 |  | 0.014 |  | 0.006 |  |
| **Tg** | **Model I** | **P** | **Model II** | **P** | **Model III** | **P** |
| VAI per IQR | 1.633 (-0.677, 3.943) | 0.166 | 0.700 (-1.689, 3.088) | 0.566 | 0.773 (-1.649, 3.196) | 0.531 |
| Quartiles of VAI |  |  |  |  |  |  |
| Q1 | Ref |  | Ref |  | Ref |  |
| Q2 | 2.624 (-3.501, 8.75) | 0.401 | 1.681 (-4.484, 7.845) | 0.593 | 1.975 (-4.209, 8.159) | 0.531 |
| Q3 | 1.302 (-4.824, 7.428) | 0.677 | 0.056 (-6.141, 6.253) | 0.986 | 0.073 (-6.157, 6.303) | 0.982 |
| Q4 | 3.815 (-2.308, 9.938) | 0.222 | 1.336 (-4.998, 7.669) | 0.679 | 1.565 (-4.869, 8.000) | 0.634 |
| *P* for trend | 0.298 |  | 0.811 |  | 0.782 |  |
| **TgAb** | **Model I** | ***P*** | **Model II** | ***P*** | **Model III** | ***P*** |
| VAI per IQR | 8.502 (4.062, 12.941) | <0.001 | 7.258 (2.678, 11.837) | 0.002 | 8.427 (3.786, 13.068) | <0.001 |
| Quartiles of VAI |  |  |  |  |  |  |
| Q1 | Ref |  | Ref |  | Ref |  |
| Q2 | 5.330 (-6.410, 17.009) | 0.374 | 3.486 (-8.323, 15.295) | 0.563 | 3.920 (-7.91, 15.75) | 0.516 |
| Q3 | 3.972 (-7.787, 15.730) | 0.508 | 1.957 (-9.915, 13.829) | 0.747 | 3.290 (-8.628, 15.219) | 0.588 |
| Q4 | 24.080 (12.32, 35.830) | <0.001 | 20.462 (8.329, 32.594) | 0.001 | 23.710 (11.399, 36.03) | <0.001 |
| *P* for trend | <0.001 |  | <0.001 |  | <0.001 |  |
| **TSH** | **Model I** | ***P*** | **Model II** | ***P*** | **Model III** | ***P*** |
| VAI per IQR | 0.208 (0.072, 0.344) | 0.003 | 0.146 (0.006, 0.287) | 0.041 | 0.149 (0.006, 0.292) | 0.041 |
| Quartiles of VAI |  |  |  |  |  |  |
| Q1 | Ref |  | Ref |  | Ref |  |
| Q2 | 0.303 (-0.057, 0.664) | 0.099 | 0.229 (-0.133, 0.592) | 0.215 | 0.235 (-0.129, 0.599) | 0.205 |
| Q3 | 0.454 (0.094, 0.815) | 0.014 | 0.364 (-0.001, 0.728) | 0.051 | 0.375 (0.009, 0.742) | 0.045 |
| Q4 | 0.717 (0.357, 1.078) | <0.001 | 0.557 (0.184, 0.929) | 0.003 | 0.579 (0.200, 0.958) | 0.003 |
| *P* for trend | <0.001 |  | 0.004 |  | 0.003 |  |
| **TPOAb** | **Model I** | ***P*** | **Model II** | ***P*** | **Model III** | ***P*** |
| VAI per IQR | 4.398 (-0.136, 8.933) | 0.057 | 2.060 (-2.611, 6.732) | 0.387 | 2.548 (-2.177, 7.273) | 0.291 |
| Quartiles of VAI |  |  |  |  |  |  |
| Q1 | Ref |  | Ref |  | Ref |  |
| Q2 | -1.85 (-13.87, 10.16) | 0.763 | -3.576 (-15.632, 8.48) | 0.561 | -3.270 (-15.33, 8.78) | 0.595 |
| Q3 | 2.530 (-9.48, 14.55) | 0.679 | 0.677 (-11.44, 12.797) | 0.913 | 1.420 (-10.720, 13.570) | 0.819 |
| Q4 | 10.54 (-1.468, 22.56) | 0.085 | 4.975 (-7.411, 17.361) | 0.431 | 6.420 (-6.120, 18.970) | 0.316 |
| *P* for trend | 0.039 |  | 0.259 |  | 0.177 |  |
| **TT3** | **Model I** | ***P*** | **Model II** | ***P*** | **Model III** | ***P*** |
| VAI per IQR | 2.357 (1.222, 3.491) | <0.001 | 3.454 (2.327, 4.582) | <0.001 | 3.641 (2.504, 4.777) | <0.001 |
| Quartiles of VAI |  |  |  |  |  |  |
| Q1 | Ref |  | Ref |  | Ref |  |
| Q2 | 3.825 (0.822, 6.828) | 0.013 | 5.960 (3.062, 8.858) | <0.001 | 6.044 (3.155, 8.933) | <0.001 |
| Q3 | 5.589 (2.587, 8.592) | <0.001 | 8.276 (5.363, 11.19) | <0.001 | 8.453 (5.543, 11.364) | <0.001 |
| Q4 | 7.391 (4.389, 10.392) | <0.001 | 10.982 (8.004, 13.959) | <0.001 | 11.480 (8.475, 14.48) | <0.001 |
| *P* for trend | <0.001 |  | <0.001 |  | <0.001 |  |
| **TT4** | **Model I** | ***P*** | **Model II** | ***P*** | **Model III** | ***P*** |
| VAI per IQR | 0.163 (0.092, 0.234) | <0.001 | 0.106 (0.033, 0.178) | 0.004 | 0.088 (0.015, 0.161) | 0.018 |
| Quartiles of VAI |  |  |  |  |  |  |
| Q1 | Ref |  | Ref |  | Ref |  |
| Q2 | 0.294 (0.107, 0.481) | 0.002 | 0.248 (0.062, 0.435) | 0.009 | 0.243 (0.058, 0.429) | 0.01 |
| Q3 | 0.412 (0.224, 0.599) | <0.001 | 0.360 (0.173, 0.547) | <0.001 | 0.357 (0.17, 0.544) | <0.001 |
| Q4 | 0.499 (0.312, 0.686) | <0.001 | 0.359 (0.168, 0.551) | <0.001 | 0.318 (0.125, 0.51) | 0.001 |
| *P* for trend | <0.001 |  | <0.001 |  | <0.001 |  |

Model I was crude model. Model II was adjusted for age, sex, edu, Marital status. Model III was adjusted for age, sex, edu, Marital status, income, drinks, Zn, Se, Na, k, smoking, hypertension, diabetes ,UIC, Cre.

